# Dynamic changes in Sox2 spatio-temporal expression direct the second cell fate decision through *Fgf4/Fgfr2* signaling in preimplantation mouse embryos

**DOI:** 10.1101/052530

**Authors:** Tapan Kumar Mistri, Wibowo Arindrarto, Wei Ping Ng, Choayang Wang, Leng Hiong Lim, Lili Sun, Ian Chambers, Thorsten Wohland, Paul Robson

## Abstract

Oct4 and Sox2 regulate the expression of target genes such as *Nanog, Fgf4* and *Utf1*, by binding to their respective regulatory motifs. Their functional cooperation is reflected in their ability to heterodimerise on adjacent *cis* regulatory elements, the composite Sox/Oct motif. Given that Oct4 and Sox2 regulate many developmental genes, a quantitative analysis of their synergistic action on different Sox/Oct motifs would yield valuable insights into the mechanisms of early embryonic development. In this study, we measured binding affinities of Oct4 and Sox2 to different Sox/Oct motifs using fluorescence correlation spectroscopy (FCS). We found that the synergistic binding interaction is driven mainly by the level of Sox2 in the case of the *Fgf4* Sox/Oct motif. Taking into account *Sox2* expression levels fluctuate more than *Oct4*, our finding provides an explanation on how Sox2 controls the segregation of the epiblast (EPI) and primitive endoderm (PE) populations within the inner cell mass (ICM) of the developing rodent blastocyst.

## Introduction

The mouse preimplantation embryo is a widely used mammalian model to study cell differentiation. Two of the earliest cell fate decisions in mammalian development take place in the preimplantation embryo. The first decision occurs at the 16-32 cell stage and sets apart the morula into two distinct lineages: the trophoblast, represented by the trophectoderm (TE) and the inner cell mass (ICM). At this stage, the TE is a single layer of epithelial cells enclosing the early blastocyst. The ICM lies at one end of the blastocyst, consisting of a pool of pluripotent cells. Later, after embryonic day 3.5 (E3.5), the ICM is further specified into the primitive endoderm (PE) and epiblast (EPI) lineages (Rossant and Tam 2009). Cells of the PE lineage subsequently differentiate into the extra-embryonic cells responsible for secreting patterning cues and providing nutrition to the developing embryo proper which consists of cells entirely from the EPI lineage.

The EPI is exclusively characterised by its *Nanog and Sox2* expression (Chambers, Colby et al. 2003; Mitsui, Tokuzawa et al. 2003; Silva, Nichols et al. 2009) while the PE is specifically characterised by *Gata4, Gata6* and *Sox17* (Morrisey, Ip et al. 1996; Morrisey, Tang et al. 1998; Koutsourakis, Langeveld et al. 1999; Aksoy, Jauch et al. 2013) while Oct4 initially persists in both (Aksoy, Jauch et al. 2013). Prior to the segregation into the PE and the EPI, the ICM shows a mosaic pattern of cells expressing *Nanog* and *Gata6* (Chazaud, Yamanaka et al. 2006). The mosaic expression of these markers does not indicate lineage specification as cells expressing the PE markers *Gata6* and *Gata4,* can be coaxed into forming the EPI lineage. The cells only become restricted to their definitive lineages at E4.5 (Chazaud, Yamanaka et al. 2006). However, studies have also shown that inner cells, which have higher *Nanog* and lower *Gata6* expression, give rise to the EPI while cells with lower levels of *Nanog* and higher levels of *Gata6* give rise to the PE (Schrode, Saiz et al. 2014; Xenopoulos, Kang et al. 2015). Therefore, it is not clear what role this difference in expression levels of lineage markers plays in the second cell fate decision of preimplantation development. In addition, how this heterogeneity emerges in the first place has also remained elusive. Studies have indicated that the *Fgf4*/*Fgfr2* signalling pathway lies upstream of this differential expression (Nichols, Zevnik et al. 1998; Guo, Huss et al. 2010; Krawchuk, Honma-Yamanaka et al. 2013). Indeed, *Fgf4* is expressed in the EPI lineage but not in the PE while *Fgfr2* is expressed in the PE but not in the EPI (Orr-Urtreger, Givol et al. 1991; Niswander and Martin 1992). Treatment with an Fgf signalling inhibitor causes the otherwise mosaic pattern of the ICM cells to generate exclusively the EPI lineage (Guo, Huss et al. 2010; Yamanaka, Lanner et al. 2010). Furthermore, both *Fgf4***-**null and *Fgfr2***-**null embryos are lethal (Feldman, Poueymirou et al. 1995; Arman, Haffner-Krausz et al. 1998). It has been further confirmed that *Fgf4* is required for the segregation of the ICM into the PE and the EPI lineages (Guo, Huss et al. 2010; Kang, Piliszek et al. 2013; Ohnishi, Huber et al. 2014). Furthermore, several studies indicate spatio-temporal differences in inner cell formation contribute to the establishment of the heterogeneity in the ICM (Pedersen 1986; Fleming 1987; Krupa, Mazur et al. 2014). Thus understanding the molecular determinants that establish this FGF4/FGFR2 signalling axis will shed light on the mechanism that established cell fate within the ICM.

In light of the current evidence from mouse preimplantation development, Sox2 emerges as a particularly interesting transcription factor to study. Along with Oct4, it has been found to regulate the expression of other genes important for preimplantation development such as *Nanog*, *Fgf4*, *Utf1*, *Pou5f1* and *Sox2* itself (Nishimoto, Fukushima et al. 1999; Ambrosetti, Schöler et al. 2000; Chew, Loh et al. 2005; Rodda, Chew et al. 2005). In the enhancers of these genes, a Sox2 binding motif, CTTTG(A/T)(A/T) (Harley, Lovell Badge et al. 1994; Wilson and Koopman 2002) is found adjacent to an octamer motif, ATGC(A/T)AA(T/A) (Verrijzer, Alkema et al. 1992) with a spacer having zero to three base pairs in between the two motifs. A recent study also enlightened the importance of an enhancer where it was illustrated that gene activation is highly correlated with the presence of an optimal motif (Farley, Olson et al. 2015). Furthermore, crystallography studies have shown that the Sox2 and Oct4 DNA binding domains heterodimerise on this motif (Reményi, Lins et al. 2003). However, unlike *Oct4*, *Sox2* levels show a dynamic pattern in the preimplantation embryo, in particular, zygotic transcription initiates within the inner cells of the morula (Guo, Huss et al. 2010). Additionally, Sox2 is known to be an activator of *Fgf4* (Yuan, Corbi et al. 1995) and a repressor of *Fgfr2* (Masui, Nakatake et al. 2007). Importantly, Sox2 is required for normal development as Sox2 null embryos fail to develop beyond early post-implantation (Avilion, Nicolis et al. 2003) and is required non-cell-autonomously via FGF4 for the development of the primitive endoderm (Wicklow, Blij et al. 2014). Collectively, these observations indicate that understanding Sox2 dynamics quantitatively is paramount to understanding the molecular mechanism of cell fate decision within the ICM.

We had previously proposed a model based on the dynamics of *Sox2*, *Fgf4*, and *Fgfr2* expression whereby the initiation of Sox2 expression in inner cells of the morula establishes the FGF signaling axis, via the up-regulation of *Fgf4* and the down-regulation of *Fgfr2*, within the ICM (Guo et al. 2010). Here we define the *cis* regulatory logic for this model by measuring the dynamic changes in Sox2 levels through preimplantation development and determining the apparent dissociation constants (aK_d_) of Sox2 and Oct4 on their respective *cis* regulatory elements on taret genes of interest. We perform these measurements through the use of fluorescent fusion proteins and fluorescent correlation spectroscopy, a single molecule sensitive fluorescence**-**based technique (Elson and Magde 1974; Jameson, Ross et al. 2009). Remarkably, our results reveal that the formation of a stable Sox2-Oct4-DNA complex on the *Fgf4* Sox/Oct motif is more dependent on the level of Sox2 than on that of Oct4. Intriguingly, the *Nanog* Sox/Oct *cis* motif does not show such a high dependency on the level of Sox2 compared to that of the *Fgf4* Sox/Oct *cis* motif. These biochemical measurements lend weight to the argument that Sox2 is indeed the driver of the earliest heterogeneity within the ICM, a heterogeneity that leads to the EPI/PrE cell fate decision.

## RESULTS

### Sox2 level increases in the ICM with time during preimplantation embryo development

Previously, it has been shown that Sox2 mRNA levels fluctuate more widely than those of Oct4 during preimplantation development (Guo, Huss et al. 2010). In order to test whether Sox2 fluctuations are also present at the protein level, we measured paternally derived zygotic GFP expression from the *Sox2* locus in *Sox2*-null embryos (Ellis, Fagan et al. 2004) following progressive cell stages during preimplantation development (Fig 1a). The earliest expression of GFP was found to be within the inner cells of the morula and later restricted to the ICM of the blastocyst. Next we measured total Sox2 levels by immunostaining (Fig 1b). While Sox2 levels are relatively high at the 4-cell stage levels decrease as development progresses to the morula. Within the blastocyst, Sox2 levels continue to recede in the TE whereas there is an increase in Sox2 from the morula to ICM (Fig 1c). This Sox2 protein dynamics closely parallels the dynamics of its mRNA level (Guo, Huss et al. 2010). The impact of this increasing concentration of Sox2 from the morula-ICM transition will be mediated through *cis* regulatory logic.

**Figure 1:**
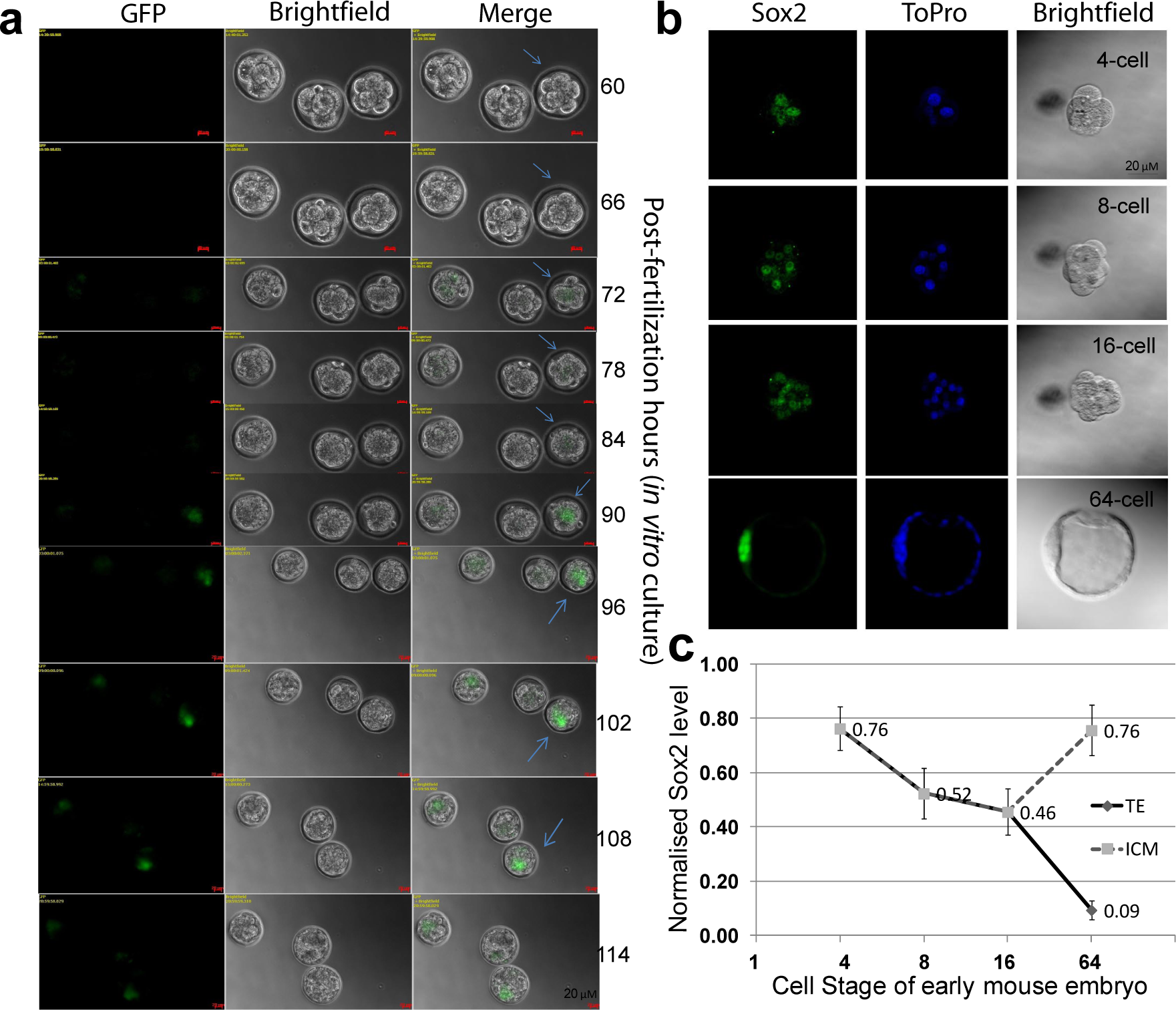
*In vivo* localisation and quantification of Sox2 level during mouse preimplantation development. (a) Microscopic observation of paternally derived zygotic *Sox2*-GFP expression in *Sox2*-null embryos (Guo, Huss et al. 2010) with the progression of time until 114 hrs from single cell zygote (guided by an arrow). GFP expression from the *Sox2* locus is observed starting from 66 hrs at 8-cell stage. The middle embryo is a wild type control. (b) Confocal Z-stack images of representative Sox2 and ToPro nuclear stained embryos. (c) Quantification of the average Sox2 level per cell in 4-16 cell embryos or in TE or ICM of 64 cell embryos shown in (b).

### Characterization of the Sox/Oct motif sequences

Despite the increasing levels of Sox2 in the nascent ICM some known target genes of Sox2 did not change (e.g. *Nanog)* while others did (e.g. *Fgf4*) (Guo et al. 2010) thus we were next interested in determining if this could be explained through specific variations in *cis* regulatory sequences mediating Sox2 binding. From a global view of Sox2 and Oct4 bound regions in embryonic stem (ES) cells it is clear that while there is sequence constraint within the sox-oct motif, there is allowable variability (Fig. 2a). This variability is in contrast to the sequence conservation seen within the particular sox-oct element in both Nanog and *Fgf4*, where sequence conservation verges on 100% identity over 100s of millions of years of cumulative evolution (Fig. 2b). Furthermore, we also observed the conservation pattern of different Sox/Oct motifs from the known genes, *Nanog, Utf1, Oct4, Sox2, and Fgf4* (Fig. 2c). Such sequence conservation strongly argues that there are functional differences between sequences that encompass the allowable sox-oct motif.

**Figure 2:**
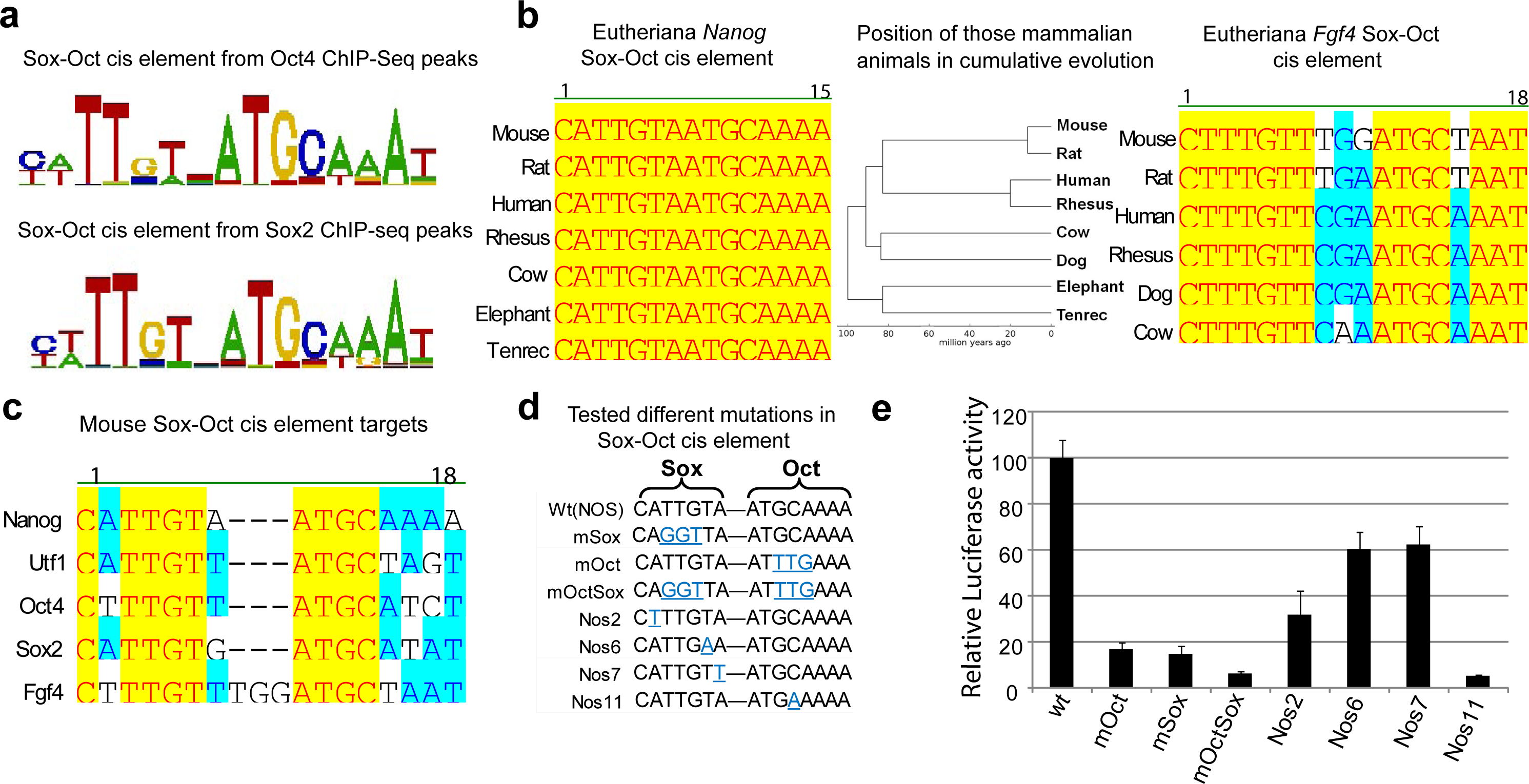
Characterisation of Sox/Oct *cis* motifs. (a) The top *de novo* sequence motifs (based on enrichment) detected by CisFinder (Sharov and Ko 2009) in Oct4 and Sox2 ChIP-Seq data from ESCs (Chen, Xu et al. 2008), shown by WebLogo. (b) Comparison of *Nanog and Fgf4* Sox/Oct *cis* motifs across different eutherian species. A phylogenetic tree illustrated the sequence conservation of those Sox/Oct motifs in cumulative evolution. (c) Alignment of the Sox/Oct motifs from the target genes *Nanog, Utf1, Oct4, Sox2* and *Fgf4*. (d) Mutations generated in the Sox/Oct motif in a 400 bp gene fragment from *Nanog* located at −289 to +117 (relative to the *Nanog* transcription start site) (Chambers 2005; Rodda, Chew et al. 2005) were subsequently tested. (e) Luciferase activity of constructs shown in (d) after transfection of F9 embryonal carcinoma cells

Through site-directed mutagenesis of the *Nanog* sox-oct motif, within the larger context of the *Nanog* 400 bp proximal promoter, we tested the functional consequences of subtle mutations on transcription as measured by luciferase activity generated in transfected F9 teratocarcinoma cells. As previously described (Rodda et al. 2005), ablation of the binding sites for Sox2 or Oct4, and in combination, through 3 bp mutations diminishes luciferase activity to below 20% of the wild-type promoter (Fig.2 d,e). When we introduced subtle single base changes of A to T, T to A, and A to T at positions 2, 6, and 7 of the sox element, respectively, there was a significant reduction in luciferase activity in all cases from 30 to 60% of wild-type levels (Fig. 2d, e). Particularly surprising was the reduction to ~60% at position 7 of the sox motif as this is the least conserved of all seven positions within the sox element (Fig. 2a, c). Thus what apparently are subtle changes to the sox-oct motif that qualitatively do not prevent binding of Sox2, have profound functional consequences on transcriptional output. We would argue that such functional consequences result in the high level of sequence conservation, through purifying selection, within specific sox-oct elements across eutherian species, particularly around these developmental control genes.

### Characterization of TFs-fluorescent fusion proteins

We hypothesized that the differential transcriptional response to increasing concentrations of Sox2 in the nascent ICM seen between *Nanog* and *Fgf4* was a result of differential binding kinetics of Sox2 and Oct4 on the associated *sox-oct* elements of these genes. As no technologies exist to measure protein-DNA binding kinetics at discrete genomic loci within living cells, we resorted to *in vitro* measurements. Our strategy was to generate quantitative measurements through Förster resonance energy transfer (FRET), FCS and EMSA on Sox2-Oct4-DNA complexes and thus required the generation of fluorescently tagged proteins.

Expression constructs were designed to produce full-length mouse Oct4 and Sox2 fused, via a four amino acid linker (GGSG), with GFP and mCherry, respectively. Initially we tested functionality of both N-terminal and C-terminal fusions. Transient transfections into mouse ES and CHO cells indicated expression of these transcription factor-fluorescent protein fusion constructs, and with nuclear localization (Fig. 3a). Western blots, using antibodies against the respective transcription factors, in nuclear lysates from ES cells transfected with these constructs further confirmed the expression of the fusion proteins (Fig. 3b). GFP-Oct4 (N-terminal) and mCherry-Sox2 appeared to be expressed at a higher level than their C-terminal tagged counterparts. Importantly, there were only two types of bands detected, namely for the fusion protein (upper bands close to 75 kDa) and the endogenous protein (lower bands in between 37 to 50 kDa).

**Figure 3:**
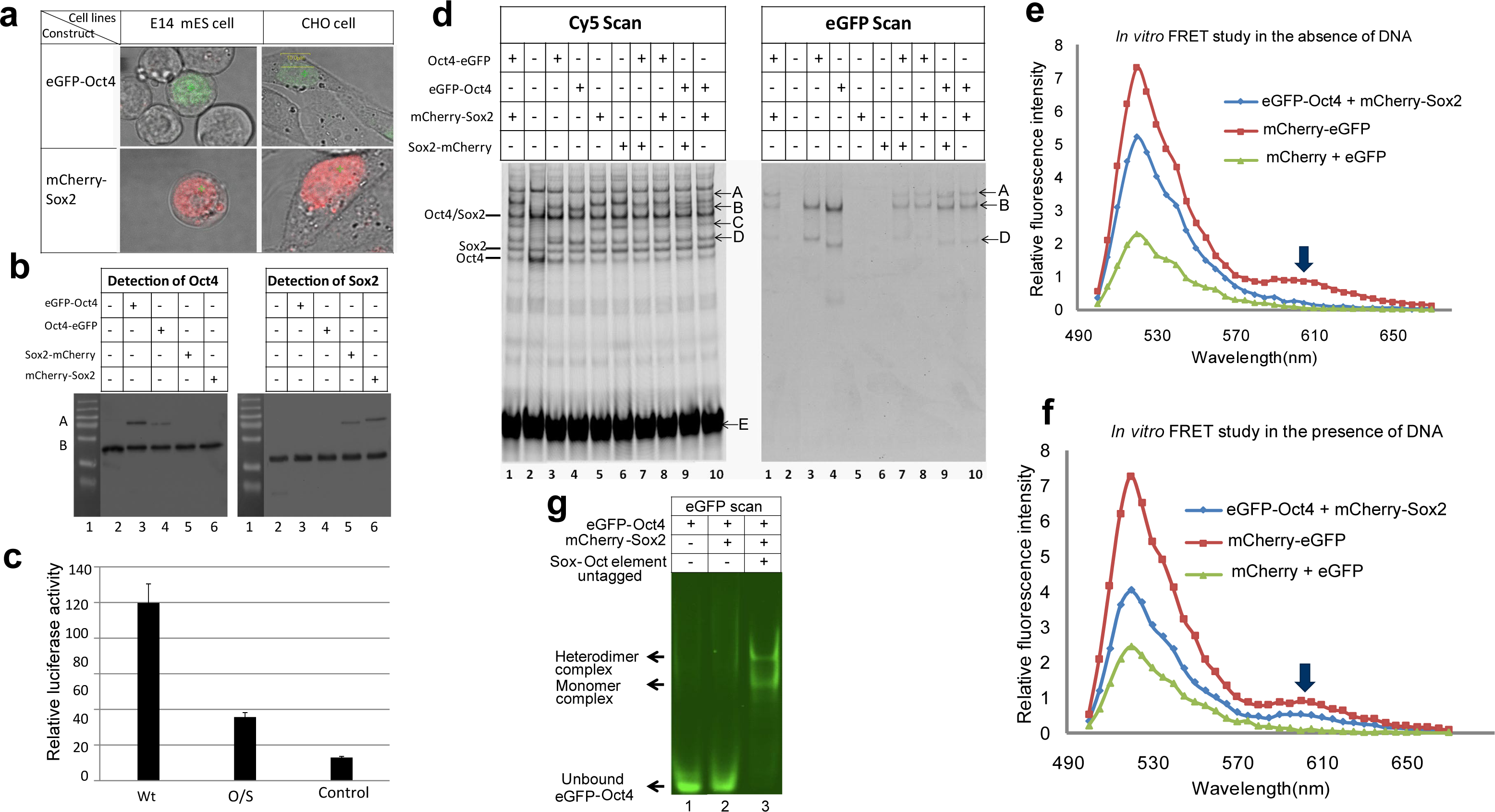
Functional determination of synthetic fusion protein constructs. (a) Confocal images of ES cells and CHO cells transfected with pCAG-GFP-Oct4-IN and pCAG-mCherry-Sox2-IN. (b) Immunoblot analysis of exogenous fusion proteins; bands A and B correspond to the fusion and wild type proteins respectively. (c) Dual luciferase assays comparing the transcriptional activity of the wild type *Nanog* Sox/Oct motif (wt) (CATTGTAATGCAAAA), a mutated *Nanog* Sox/Oct motif (O/S) (CA<u>GGT</u>TAAT<u>TTG</u>AAA), and a *Nanog* promoter with a deletion form X – y that removes ONLY the Sox/Oct motif (control). (d) DNA binding of fusion proteins (GFP scan) compared to wild type proteins (Cy5 scan) by EMSA from ES cells nuclear lysates. A – heterodimer complex (GFP-Oct4/mCherry-Sox2/NSO element); B – heterodimer complex (GFP-Oct4/wild type Sox2/NSO element); C – monomer complex (mCherry-Sox2/NSO element); D – monomer complex (GFP-Oct4/NSO element). (e-f) Emission intensity collected at >500 nm wavelength upon GFP excitation by 488nm laser. FRET analysis was performed on nuclear extracts isolated from transfected CHO cells in the absence (e) or the presence (f) of DNA containing the *Nanog* Sox/Oct motif. The arrow indicates the emission maxima of mCherry. (g) Complex formation shown by FP-EMSA detecting GFP-Oct4 in the presence or absence of unlabelled DNA and mCherry-Sox2.

We next tested our fusion constructs for their ability to activate transcription in a luciferase promoter assay. CHO cells were used in these experiments as they lack endogenous Oct4 and Sox2 (Fig. S1a). In transient transfection assays, co-transfection of the Oct4 and Sox2 fusion constructs with the wild type *Nanog* promoter (Wt) resulted in significant luciferase activity above that of the control that lacked the promoter and the mutated promoter (O/S) where mutations applied to the Sox/Oct motif (Fig. 3c), indicating that the fusion constructs have the ability to drive transcription from the *Nanog* promoter.

We further sought to determine whether these modified TFs retained the ability to bind DNA similarly to their endogenous counterparts. We tested, by EMSA, nuclear lysates from ES cells transfected with these plasmid constructs for their ability to bind an oligonucleotide containing the *Nanog* Sox/Oct motif. We observed that both endogenous proteins and their fusion counterparts are capable of binding this DNA element as monomers and heterodimers (Fig. 3d). Differentially shifted bands indicate the fusion proteins can heterodimerise with both their respective endogenous and fusion protein partners. The fusion containing complexes are readily detectable despite these proteins being expressed at lower levels (Fig. 3b) and requiring to compete with their endogenous counterparts in this assay. These results indicate these fusion proteins are as competent as their wild type proteins in binding to DNA containing the Sox/Oct motif. Finally, we confirmed the physiological function of the N-terminal fusion proteins, GFP-Oct4 and mCherry-Sox2 for their ability to rescue ES cells in which the corresponding endogenous TF alleles had been deleted (Mistri, Devasia et al. 2015).

### The Sox2-Oct4 protein-protein interaction requires DNA

Having demonstrated that the Oct4 and Sox2 fusion proteins perform functionally similarly to their endogenous counterparts, we next sought to utilize GFP-Oct4 and mCherry-Sox2 to quantify their combinatorial binding interplay on the Sox/Oct motif. Utilizing FRET we quantitatively investigated the formation of the Sox2 and Oct4 heterodimer complex with the *Nanog* Sox/Oct motif in solution using nuclear extracts from transfected CHO cells. We examined Sox2-Oct4 interactions in the presence and absence of DNA to understand the DNA dependency of Sox2-Oct4 complex formation. No FRET signal was observed from a solution containing GFP-Oct4 and mCherry-Sox2 in the absence of DNA (Fig. 3e) however, when DNA containing a *Nanog* Sox/Oct motif was included a distinct FRET signal was detected (Fig. 3f). This observation indicates that the DNA brings GFP-Oct4 and mCherry-Sox2 into close proximity enabling successful energy transfer from GFP to mCherry, through binding to the Sox/Oct motif. In further validation our FP-EMSA assay also did not detect Sox2-Oct4 interaction unless sox-oct DNA was present (Fig. 3g) nor are any multimers of GFP-Oct4 detected. These results indicate that heterodimer complexes are only possible in the presence of DNA which is in agreement with our previous work (Chen, Tapan et al. 2012; Mistri, Devasia et al. 2015).

### Determination of apparent dissociation constants (aK_d_) by FCS and EMSA

Understanding *Fgf4* gene regulation requires the quantitative measurement of mCherry-Sox2 binding to the *Fgf4* Sox/Oct motif compared to that with the *Nanog* and *Utf1* motifs. For these quantitative studies we used FCS and EMSA as complementary methods. While FCS has been used to measure aK_d_s in lysate as well as in live cells and zebrafish embryos (Shi, Foo et al. 2009; Mistri, Devasia et al. 2015), FP-EMSA was applied for quantitative aK_d_ measurements for full-length fusion proteins as one can measure the concentration of fusion protein by FCS even in unpurified nuclear lysate (Mistri, Devasia et al. 2015). The titration strategy is shown in Fig. 4a. Direct evidence of complex formation on the *Fgf4* Sox/Oct motif under different titration conditions was compared between EMSA generated gel images and FCS generated ACF curves (Fig. 4b). In the presence of 72 nM mCherry-Sox2, a titration of GFP-Oct4 to the *Fgf4* Sox/Oct motif yielded aK_d_s of 25.2 ± 4.1 nM and 25.3 ± 2.2 nM from EMSA and FCS, respectively. (Fig. 4c). On the other hand, a titration of mChery-Sox2 to the same *Fgf4* Sox/Oct motif in presence of 40 nM GFP-Oct4 produced aK_d_ of 23.2 ± 1.2 nM and aK_d_ of 24.0 ± 3.0 nM from EMSA and FCS, respectively (Fig. 4d). Notably, both FCS and EMSA provided similar values within the margins of standard deviation, strengthening the reliability of our quantitative findings on TF-DNA binding interactions.

**Figure 4:**
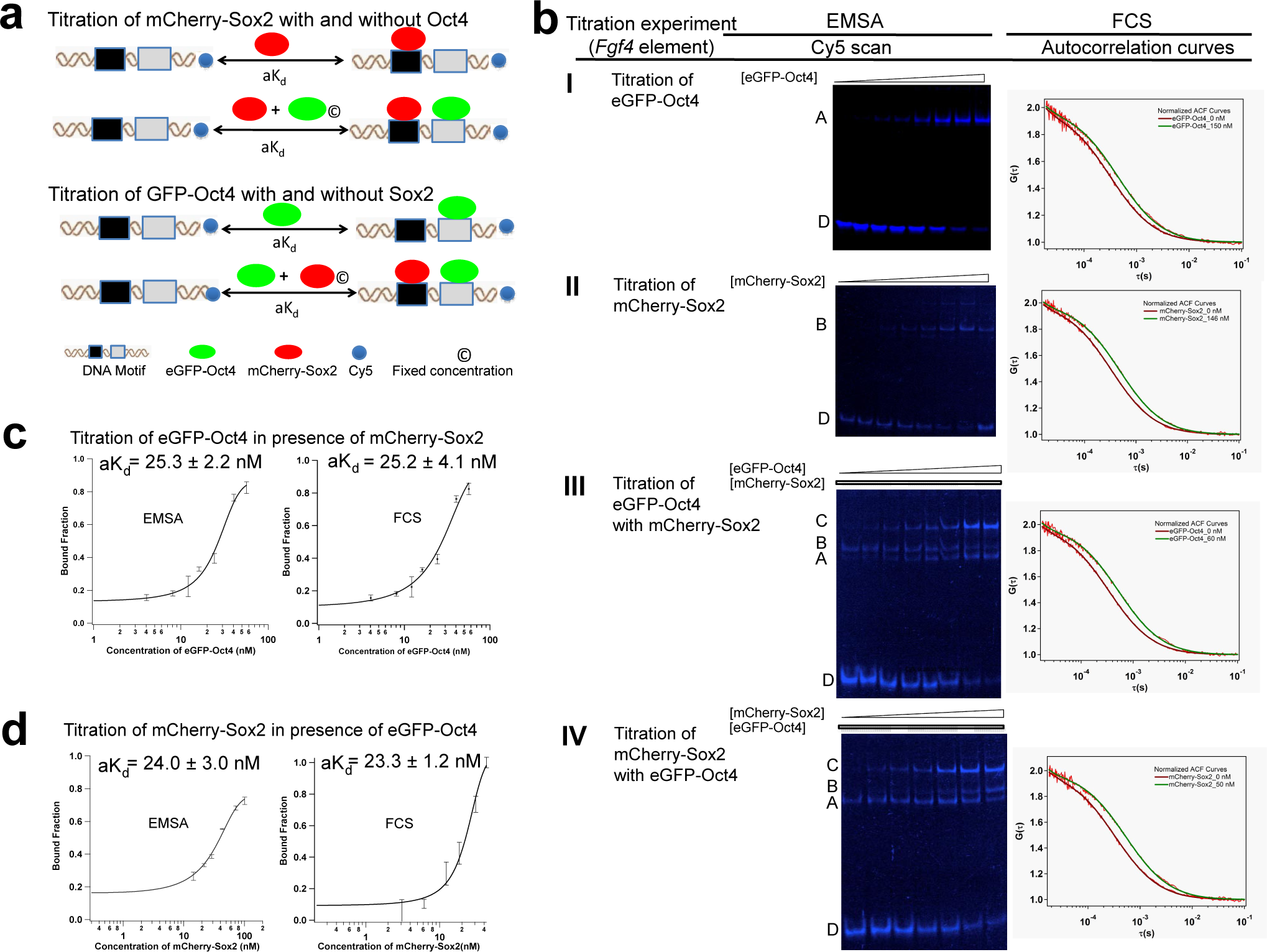
Comparison of FCS and EMSA derived aK_d_. (a) Conceptual scheme showing the titration of GFP-Oct4 in the absence and presence of mCherry-Sox2 and vice versa. (b) EMSA scans and autocorrelation curves generated from FCS assays for the *Fgf4* Sox/Oct motif titrating with (I) GFP-Oct4; (II) mCherry-Sox2; (III) GFP-Oct4 in the presence of a fixed mCherry-Sox2 concentration and (IV) mCherry-Sox2 in the presence of a fixed GFP- Oct4 concentration. A: monomer (GFP-Oct4/DNA); B: monomer (mCherry-Sox2/DNA); C: heterodimer (GFP-Oct4/mCherry-Sox2/DNA); D: free Cy5-DNA. (c-d). Comparison of the bound fraction vs. total protein concentration plots with the *Fgf4* Sox/Oct motif obtained by EMSA (left panel) or FCS (right panel).

### Influence of sequence variation within the Sox/Oct motif on protein-DNA binding affinity

Previously we had observed that sequence changes in the conserved and non-conserved regions of the Sox2 motif had an impact on transcriptional activity (Fig. 2E). Such activity may be linked to the degree of Oct4 and Sox2 binding affinities for these DNA elements and consequently impact the formation of a stable heterodimer. The change in the sequence of the Sox2 binding site “CATTGT<u>A</u>” in *Nanog* to “CATTGT<u>T</u>” in *Utf1* revealed a slight decrease in the *Sox2* binding affinity as the aK_d_ increased to 44.0 ± 9.8 nM from 31.7 ± 4.6 nM as measured by FCS. The change in the 7^th^ position of the sequence “CATTGT<u>A</u>” in *Nanog* to “CATTGT<u>G</u>” in *Sox2* revealed a slight decrease in the *Sox2* binding affinity as the aK_d_ increased to 66.1 ± 18.2 nM from 44.0 ± 9.8 nM. Additionally we observed a slight decrease in the *Sox2* binding affinity when both 2^nd^ and 7^th^ positions were changed to “C<u>T</u>TTGT<u>T</u>” in *Fgf4* from “C<u>A</u>TTGT<u>A</u>” in *Nanog* corresponding to an increase to ~ 70 nM for the aK_d_ (Fig S3 and Table 1). Our result demonstrates that the variable positions in the heptamer sequence play an important role in the binding interactions of Sox2 with the Sox/Oct motif while the conserved positions are key for strong interactions. The variable positions in the Sox2 binding sequence (CtTTGTt) of different Sox/Oct motifs create diversity in DNA binding affinities of Sox2.

**Table 1:**
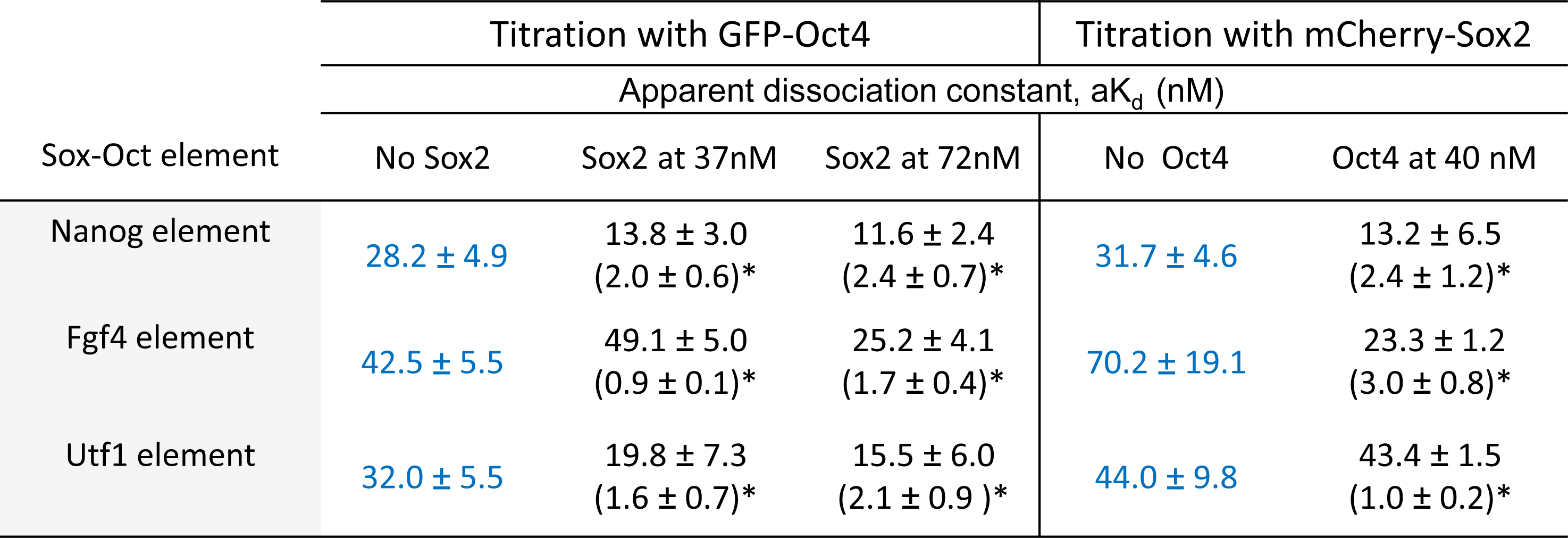
(a) Synergism correlates with the Sox2 concentration. aK_d_ measurements from different titrations of GFO-Oct4 against *Nanog, Fgf4,* and *Utf1* elements in the presence and absence of mCherry-Sox2 by FCS and vice versa.

| Sox-Oct element | Titration with GFP-Oct4 |  |  | Titration with mCherry-Sox2 |  |
| --- | --- | --- | --- | --- | --- |
|  | Apparent dissociation constant, aK<sub>d</sub> (nM) |  |  |  |  |
|  | No Sox2 | Sox2 at 37nM | Sox2 at 72nM | No Oct4 | Oct4 at 40 nM |
| Nanog element | 28.2 ± 4.9 | 13.8 ± 3.0<br>(2.0 ± 0.6)* | 11.6 ± 2.4<br>(2.4 ± 0.7)* | 31.7 ± 4.6 | 13.2 ± 6.5<br>(2.4 ± 1.2)* |
| Fgf4 element | 42.5 ± 5.5 | 49.1 ± 5.0<br>(0.9 ± 0.1)* | 25.2 ± 4.1<br>(1.7 ± 0.4)* | 70.2 ± 19.1 | 23.3 ± 1.2<br>(3.0 ± 0.8)* |
| Utf1 element | 32.0 ± 5.5 | 19.8 ± 7.3<br>(1.6 ± 0.7)* | 15.5 ± 6.0<br>(2.1 ± 0.9 )* | 44.0 ± 9.8 | 43.4 ± 1.5<br>(1.0 ± 0.2)* |

When we consider the Oct4 motif in terms of its binding affinity, we observed that the first four base pairs, ATGC, in the Octamer motif are conserved throughout the five genes. From our luciferase assay (Fig. 2c) we know that this conserved region has an important role in Oct4 binding to its motif; but the non-conserved region (5^th^ to 8^th^ position in the octamer motif) also has an important role in the degree of binding affinity as we have seen from the aK_d_ values determined on different Sox/Oct motifs. Comparing between the *Oct4* and the *Sox2* Sox/Oct motifs, we found that the 7^th^ position plays a role in increasing the aK_d_ values from 7.7 ± 1.1 nM to 15.9 ± 1.6 nM. We also looked into the *Utf1* and *Fgf4* motifs and observed that the 7^th^ position plays an important role in increasing the aK_d_ value from 32.0 ± 5.5 nM to 42.5 ± 5.5 nM. The 5^th^ and 6^th^ position displayed a dramatic change when “AT” was replaced by “TA”; the value increased to 25 nM, as compared to the *Sox2* and *Fgf4* Sox2 binding site (Fig S3 and Table 1). Therefore, our comparative aK_d_ measurements demonstrate the role of variable positions of different Sox/Oct motifs in influencing binding affinity.

Interestingly, we observed that ATGC is highly conserved and and thus could be anticipated that ATGC has more influence on Oct4 binding specificity to the octamer sequence of Sox/Oct motifs. Therefore, in addition to FCS affinity measurements, we performed a FP-EMSA with GFP-Oct4 on differently mutated motifs (see supplementary information) where we applied mutations in the octamer motif sequence (ATGCAAAA). The results showed that Oct4 binding affinity is more strongly correlated with the first 4 base pairs of the octamer motif sequence (ATGC) than the last 4 base pairs (AAAA) (Fig. S4a). We further attempted to understand the effective influence of a single base pair compared to the collective influence from the conserved base pairs of the octamer sequence (ATGC). In our FP-EMSA experiment, we noticed that the sequence AT has a stronger influence than GC (Fig. S4b). These results indicate the Sox/Oct motif with a mutation in the ATGC region would be less potent in binding with Oct4.

### The role of Sox2 concentration on its synergistic interaction with Oct4

Having now established that DNA is necessary for the formation of a stable complex between GFP-Oct4 and mCherry-Sox2, we further investigated the importance of protein level for the formation of a stable ternary complex. We measured the *in vitro* aK_d_ for mCherry-Sox2 and GFP-Oct4 independently and when in solution together by FCS in two separate titrations (Fig. 4a). The main objective was to evaluate whether the ternary complex on the *Fgf4* Sox/Oct motif shows any significant response to the level of mCherry-Sox2. We noticed that in the presence of mCherry-Sox2 as a cofactor, GFP-Oct4 showed a higher affinity for the DNA, thus driving exclusive heterodimer formation. Similarly, the presence of GFP-Oct4 as a cofactor aided the binding of Sox2 to these Sox/Oct motifs thus providing evidence that Oct4 and Sox2 have synergistic effects for *Nanog, Fgf4 and Utf1* (Table 1). However, we noticed a significant difference in stable ternary complex recruitment among the *Fgf4*, *Nanog* and *Utf1* Sox/Oct motifs whereby the *Fgf4* Sox/Oct motif needs higher levels of Sox2 rather than of Oct4 for the formation of a stable Oct4-Sox2-DNA complex.

The enhanced binding of Oct4 to *Nanog, Fgf4* and *Utf1* depends on the concentration of Sox2 as validated by the increase in the apparent cooperativity factor at the higher Sox2 cofactor concentration (Table 1). The individual titration of the *Fgf4* Sox/Oct motif with Sox2 gave an aK_d_ of 70.2 ± 19.1 nM (Table 1). Due to the lower binding affinity of Sox2 to the *Fgf4* motif as compared to that of Nanog as well as to that of *Utf1*, the influence of Sox2 on Oct4 binding is smaller, giving an apparent cooperativity factor close to 1 at lower Sox2 concentration of 40 nM. At a higher Sox2 concentration of 72 nM, a greater apparent cooperativity factor of 1.7 ± 0.4 was obtained (Table 1). In contrast, *Nanog* and *Utf1* showed a similar kind of synergistic effect at a lower Sox2 concentration of 32 nM. This suggests that a higher Sox2 concentration is essential for increasing the binding affinity of Oct4 for the *Fgf4* Sox/Oct motif. Therefore, from our *in vitro* titration data we conclude that *Fgf4* motif needs high levels of Sox2. It has also been reported that *Sox2* and *Fgf4* correlate with each other at the mRNA level during the preimplantation development (Guo et al. 2010). Consistent with this observation, we too found that the protein levels of Sox2 showed similar trend as seen earlier (Fig. 1). Therefore, it will be interesting to address further whether the expression of *Fgf4* depends on the level of Sox2 in vivo. We sought the answer in Sox2-null embryos by RT-qPCR.

### Comparison of Sox2 binding affinity between *Fgf4* and *Fgfr2 cis* regulatory motifs

It has been previously hypothesized that the *Fgfr2* gene is a direct target for Sox2 (Chen, Xu et al. 2008). We analysed the published ChIP-Seq data for Sox2 in ES cells and identified a Sox *cis* motif (Fig. 5a and Fig. S2). We further looked for the sequence conservation of the novel Sox *cis* motif which illustrated that the motif is only conserved in rodents but with one base pair mismatch in canines, bovines, elephants and opossums. However, it is widely variable for humans. This suggests that this element is important for rodents but not for humans (Roode, Blair et al. 2012). We next sought to verify whether Sox2 directly binds to the novel motif as it does to the *Fgf4* Sox/Oct motif, as Sox2 was described to be a regulator of earlier *Fgf4* and *Fgfr2* genes expression (Guo, Huss et al. 2010) Furthermore, Masui et al., 2007 had experimentally shown that the absence of Sox2 favours *Fgfr2* upregulation which suggests that Sox2 may be working as a repressor of *Fgfr2* (Masui, Nakatake et al. 2007). We performed an EMSA assay using mCherry-Sox2 in CHO nuclear cell lysate with the *Fgfr2* and *Fgf4* motifs. We noticed that mCherry-Sox2 formed a stable monomer with both DNA elements (Fig. 5b). We further quantified the binding affinities of Sox2 to these motifs by FCS (Fig 5c&d). Our result showed that both *Fgfr2* (aK_d_ values is 81.2 ± 15.1 nM) and *Fgf4* (aK_d_ values is 70.2 ± 19.1 nM) require high concentrations of Sox2 for stable complex formation (Fig. 5c&d). However, the presence of the Oct4 binding motif in the *Fgf4* Sox/Oct motif favours stable complex formation even at low concentrations of Sox2 (Table 2). This could be the reason for the good correlation between *Fgf4* expression and Sox2 level during preimplantation development. On the other hand, *Fgfr2* shows minimal expression where the Sox2 level is high such as in the EPI and the opposite happens in the TE lineage. This suggests that Sox2 works as an activator of *Fgf4* and repressor of *Fgfr2.* Therefore, it will be interesting to address further whether the expression of *Fgf4* depends on the level of Sox2 *in vivo*. We sought the answer in Sox2-null embryos by RT-qPCR.

**Figure 5:**
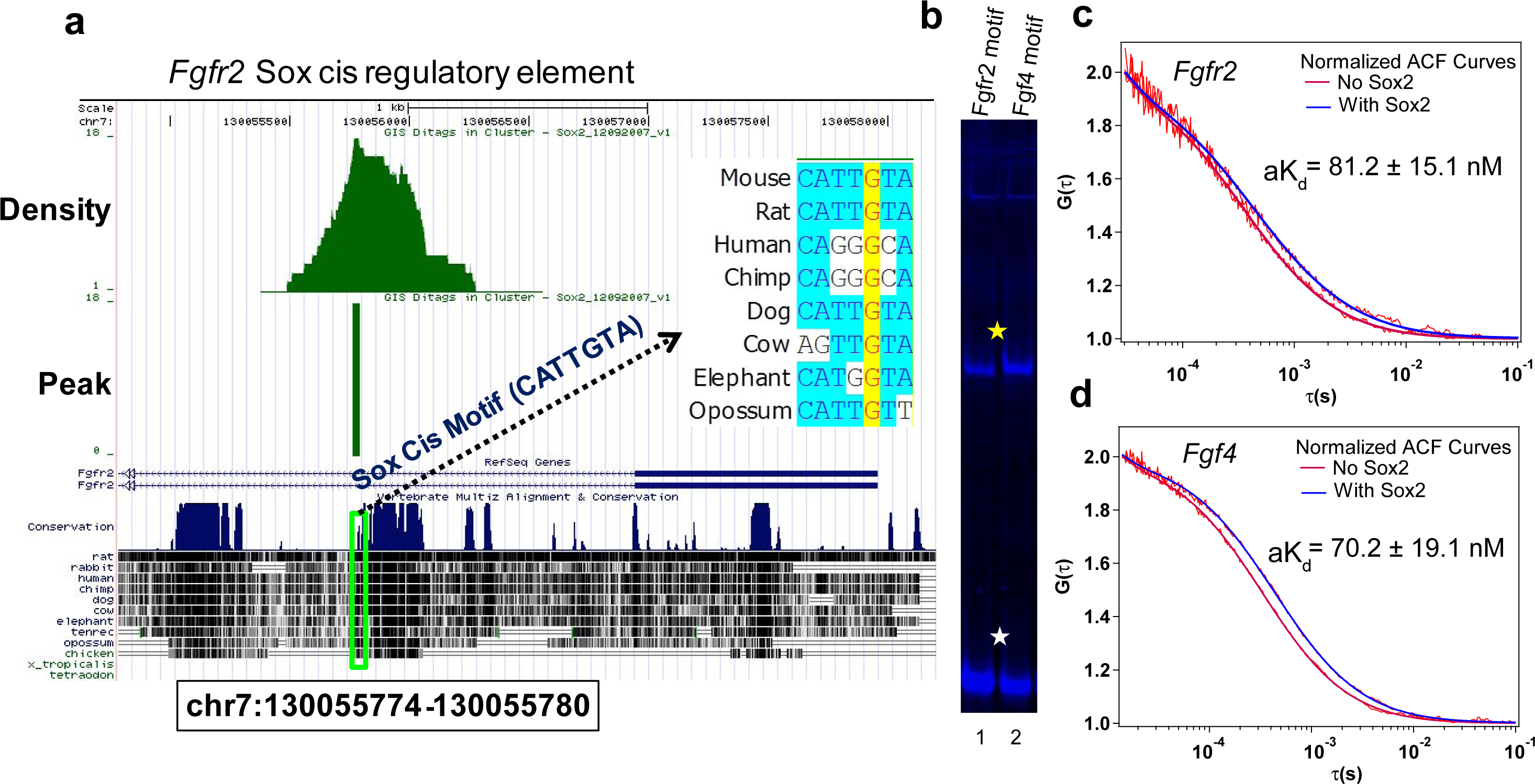
Direct binding of Sox2 protein to *cis* elements of *Fgf4* and *Fgfr2*. (a) Identification of the Sox *cis* regulatory element near the transcription start site of *Fgfr2* from Sox2 ChIP-Seq data in ES cells (Chen, Xu et al. 2008). Sequence conservation of the novel motif across different mammalian species is shown. (b) EMSA comparison of the binding of Sox2 to the novel Sox *cis* motif and the *Fgf4* Sox/Oct *cis* motif; lane 1: *Fgfr2* element, Lane 2: *Fgf4* element. Yellow and white stars indicate Sox2-DNA complex and free DNA respectively. (c-d) Two representative normalized autocorrelation (ACF) curves at no Sox2 as well as at high level of Sox2 shown for *Fgfr2* (c) and *Fgf4* (d) elements. The measured aK_d_s from each titration assay measured by FCS technique were compared.

### Validation of the role of Sox2 on its target genes in Sox2-null embryos by RT-qPCR

Taken together, our present and published work (Guo, Huss et al. 2010) suggests that Sox2 works as regulator for both *Fgf4* (positively) and *Fgfr2* (negatively). To investigate this further, we performed RT-qPCR in total mRNA samples derived from Sox2-null embryos (Ellis, Fagan et al. 2004). We noticed that the expression of *Fgf4* is minimal compared to that of the housekeeping gene, Actin β On the other hand, expression of *Fgfr2* is higher than that of Actin β(Fig. 6b). We also observed that expression of *Nanog* is not influenced by the absence of Sox2 which further supports our *in vitro* data where we observed that the *Nanog* Sox/Oct motif is capable of making stable ternary complexes at lower Sox2 concentrations than the *Fgf4* Sox/Oct motif. It should be noted that maternal Sox2 is still present in early blastocyst and this could be sufficient for the expression of *Nanog* but not for that *Fgf4* and *Fgfr2* as they need higher Sox2 levels (Avilion, Nicolis et al. 2003). RNA-Seq data further support our findings as the expression of *Fgf4*’s was minimal in Sox2-null embryo in comparison with wild type embryos (data not shown).

**Figure 6:**
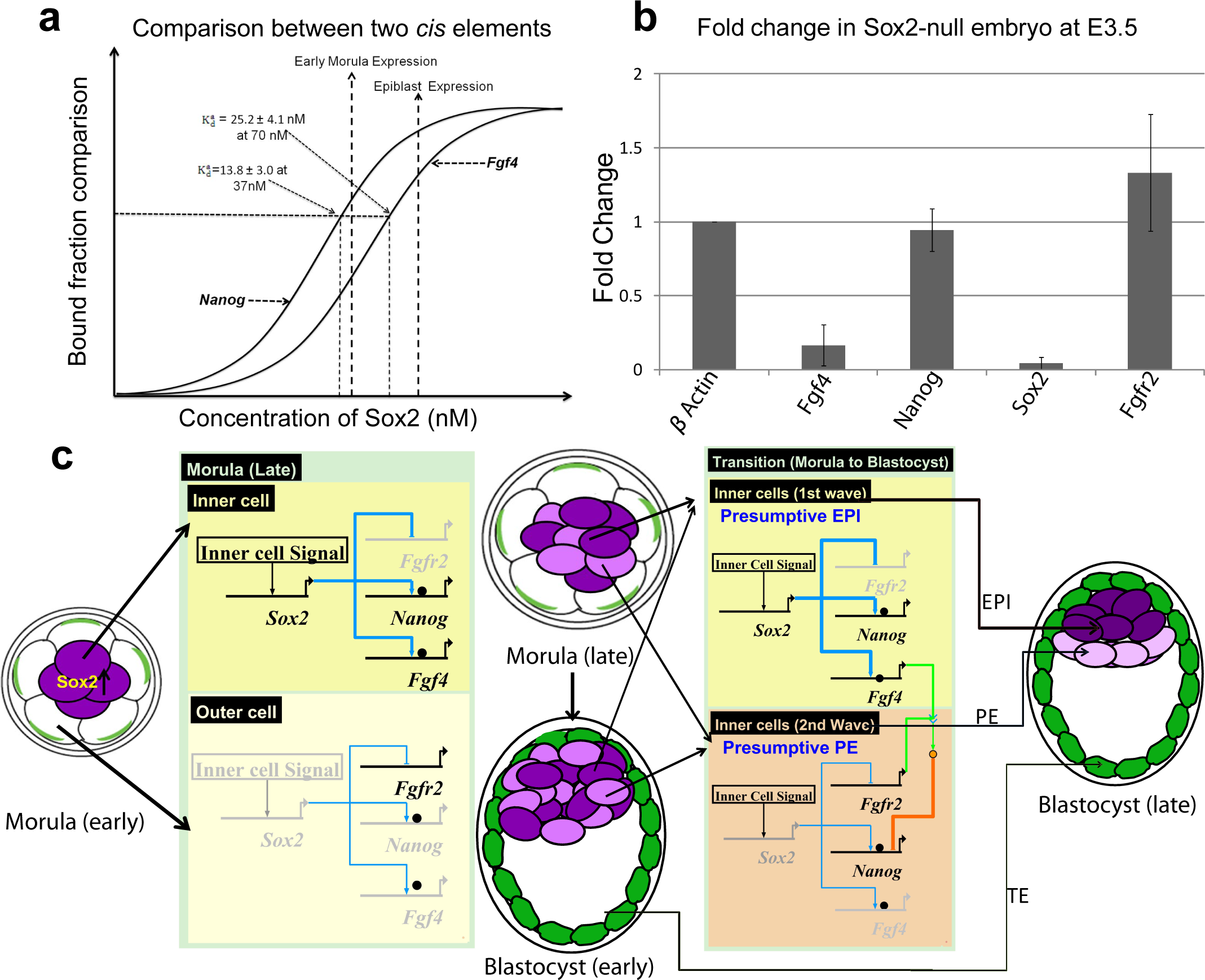
Bridging in *vitro* measurements with *in vivo* Sox2 levels of early embryos. (a) Hypothetical diagram showing relation of aK_d_ to gene expression with respect to Sox2 concentration is depicted. (b) Relative mRNA levels of *Fgf4*, *Nanog,* and *Fgfr2* in *Sox2*-null embryos at E3.5 were compared to that of WT embryos by RT-qPCR. (c) Gene regulation model for *Fgf4, Nanog* and *Fgfr2* during the segregation of ICM into PE and EPI by Bio Tapestry software (Longabaugh 2012). The thickness of the blue lines refers to the Sox2 level; the black ball indicates Oct4. Deep and light purple refer to the early inner cells and the late inner cells respectively. Green shade refers to trophectodermal lineage.

## Discussion

Whether the segregation of ICM into PE and EPI is driven by stochastic or deterministic events is currently controversial (Chazaud, Yamanaka et al. 2006; Morris, Teo et al. 2010). However, this differentiation event depends on Fgf/Erk signaling which in turn depends upon communication between inner cells expressing *Fgf4* and inner cells expressing *Fgfr2* (Chazaud, Yamanaka et al. 2006; Guo, Huss et al. 2010; Kang, Piliszek et al. 2013; Ohnishi, Huber et al. 2014). In this study, we provide evidence in support of a model in which temporal alterations in the Sox2 concentration differentially regulate expression of *Fgf4* and *Fgfr2, thereby driving* segregation of ICM into PE and EPI.

To assess key differences in the protein-DNA binding interactions in the light of what is known about the second differentiation event of mouse embryonic development; we measured the binding kinetics of Oct4 and Sox2 to different Sox/Oct enhancers specific to *Nanog*, *Fgf4* and *Utf1*. Our *in vitro* protein-DNA binding analyses indicate that stable protein-DNA complex formation is dependent not only on the DNA sequence specificity but also on the concentration of proteins involved. Interestingly, binding of Sox2 to the *Fgf4* Sox/Oct motif requires a higher concentration of Sox2 (~ 2 fold) than is needed for similar complex formation on Sox/Oct motifs from *Nanog* or *Utf1*. Direct binding of Sox2 to the *Fgfr2* and *Fgf4* Sox/Oct motifs and the requirement of a high Sox2 concentration for formation of a stable Sox2/DNA complex further lead us to question whether the expression of *Fgf4/Fgfr2* correlates with the high level of Sox2 *in vivo*. In Sox2-null embryos at E3.5 (and therefore, in the absence of paternal Sox2), *Fgf4* is down-regulated and *Fgfr2* is up regulated. These findings argue that *Fgf4/Fgfr2* expression is highly dependent on the Sox2 concentration.

Based on our *in vitro* and *in vivo* results, we propose a model considering *Fgf4* and *Nanog* which adds more clarity to the second cell fate decision during mouse development (Fig. 6a). At 37 nM Sox2, the aK_d_ value for the *Nanog* Sox/Oct motif is 13.8 ± 3.0 nM with an apparent cooperativity factor of 2.0 ± 0.6 (Table 1), while the aK_d_ for the *Fgf4* Sox/Oct motif is 42.5 ± 5.5 nM with an apparent cooperativity factor of 0.9 ± 0.1 (Table 1). Therefore, at low Sox2 concentration, such as 37nM, *Nanog* may be expressed but *Fgf4* may not be expressed. This agrees with mRNA expression analysis (Fig. 6b) at E3.5, showing that *Nanog* is expressed but *Fgf4* is not. At 72 nM Sox2, the aK_d_ value and apparent cooperativity factor for the *Fgf4* Sox/Oct motif was 25.2 ± 4.1 nM and 1.7 ± 0.4 respectively, whereas the aK_d_ and the apparent cooperativity factor for the *Nanog* motif were relatively unchanged. At this elevated Sox2 concentration both *Nanog* and *Fgf4* may be expressed.

From the above argument, we propose the gene regulation model illustrated (Fig. 6c) controlling the segregation of the ICM into the EPI and the PE. As the zygotic Sox2 expression is detectable in morula, we consider development from the morula to the late blastocyst. The morula, 16-cell stage embryo, consists of a group of inner cells surrounded by outer cells. In the inner cells, the zygotic expression of Sox2 starts but initially at a concentration not high enough to drive upregulation of *Fgf4* and downregulation of *Fgfr2.* In contrast, expression of Sox2 at this stage is absent in the outer cells and the maternal Sox2 protein level is depleting, resulting in downregulation of *Fgf4* and upregulation of *Fgfr2*. This scenario becomes more critical for the outer cells at the end of another round of cell division where the Sox2 level decreases further resulting in no expression of *Fgf4* and *Nanog*; but the clear expression of *Fgfr2*. At the end of the 32 cell stage, those outer cells behave like presumptive trophectoderm (TE). Prior to the cavitation process, there remains heterogeneity among inner cells, due to a second wave of inner cell formation after cell division of the 16-cell morula. At the end of the 32-cell stage, the early inner cells (deep purple) will already be expressing *Fgf4* and downregulate *Fgfr2* whereas the late inner cells (light purple) have high levels of *Fgfr2*. These considerations are consistent with a previous report that also suggested that cells generated in the second wave express higher levels of *Fgfr2* than those from the first wave (Morris, Graham et al. 2013).

After cavitation, the late morula proceeds to the early blastocyst stage where the outer cells are already fated to become TE and the mixed population of early and late inner cells create the ICM. We assigned to the early inner cells in the ICM as the label of presumptive EPI and to the late inner cells that of presumptive PE. At this stage of embryo development, the early inner cells possess a high level of *Fgf4* whereas the late inner cells possess a high level of *Fgfr2*. Therefore, the late inner cells come in direct contact with the signalling output of *Fgf4*-expressing early inner cells, resulting in the downregulation of a number of pluripotency genes including, *Nanog, Klf2,* and *Esrrb* in the late inner cells and upregulation of PE-specific genes including *Gata6, Gata4,* and *Sox17.* Therefore, the late inner cells go on to become the PE lineage. In contrast, the early inner cells maintain the upregulation of the pluripotency genes and increased zygotic Sox2 levels replenish the depleting level of maternal Sox2 protein. Consequently, they commit to the EPI lineage. In summary, the *in vitro* protein-DNA binding data and *in vivo* analysis of Sox2 levels controlling the expression of *Fgf4* and *Fgfr2*, allows us to argue that Sox2 increasing levels in the inner cells could be a determinant for the segregation of the ICM into the PE and the EPI cell lineages.

## Materials And Methods

### Electrophoretic mobility shift assay (EMSA)

EMSA was carried out as described (Rodda, Chew et al. 2005; Chen, Tapan et al. 2012). Details of quantitative titration assay are provided in supplementary information. Additionally a fluorescence protein based EMSA (FP-EMSA) was also performed as described (Chen, Tapan et al. 2012; Hutchins, Choo et al. 2013). All the oligonucleotides used for EMSA are provided in the supplementary information.

### Luciferase Assay

CHO cells were cultured in Dulbecco’s modified Eagle’s medium with high glucose (Invitrogen), 10% standard fetal bovine serum (Hyclone), and 1% penicillin/streptomycin and maintained at 37°C with 5% CO_2_. For a 24-well plate, 0.5 μg of EO plasmid, 0.5 μg of MS plasmid and 0.3 μg of pGL3 NSO plasmid were co-transfected per well using Lipofectamine 2000 (Invitrogen) according to the manufacturer’s instructions. 0.05 μg of *Renilla* luciferase plasmid (pRL-TK from Promega) was co-transfected as an internal control. Firefly and *Renilla* luciferase activities were measured 24 hours post transfection using the Dual Luciferase Kit (Promega) and a Centro LB960 96-well luminometer (Berthold Technologies). Alternatively, F9 embryonal carcinoma (EC) cells were used in the luciferase assay for motif characterization to understand the importance of variable positions in the Sox/Oct motif sequence. Cell culture, transfection and sample preparation for F9 EC cells were performed as described earlier (Rodda, Chew et al. 2005).

### Concentration measurement of fusion protein

The concentration of fusion proteins in unpurified nuclear lysate by fluorescence correlation spectroscopy (FCS) was measured as described (Buschmann, Krämer et al. 2009; Mistri, Devasia et al. 2015). See the supplementary section for the theoretical models used in FCS analysis.

### Data analysis (EMSA& FCS) for aK_d_ determination

The apparent dissociation constants,aK_d_, were determined as described earlier (Mistri, Devasia et al. 2015).

### Immunocytochemical Staining and Image J based semi-quantification

Embryos were fixed in 2.5% PFA for 15 min at 37°C, washed with Triton (0.1% in PBS; 5 min), Triton (0.5% in PBS; 20min), Triton (0.1% in PBS; 5 min), and BSA/Tween (0.1%BSA and 0.01% Tween in PBS; 30 min). After incubation with 1° antibody (Sox2-Y17) in BSA/Tween (60 min), embryos were washed with BSA/Tween (3x15min), and then incubated with 2° antibody (Goat anti-Rabbit IgG) conjugated with Alexa Fluor 488 (Molecular Probes, Carlsbad, CA) for an additional hour. Following BSA/Tween washes (3x15min), embryos were passed through increasing concentrations of mounting solution containing To-Pro prior to final mounting. Images were captured with a confocal microscope (LSM 510 META; Zeiss, Thornwood, NJ). All the Z-stack images from an individual embryo were grouped into one stack picture based on average fluorescence intensity employing image J software (NIH). Nuclear staining dye To-Pro was used as a control for normalizing the fluorescence intensity of Sox2 targeted antibody tagged with Alexa Fluor 488. It is known that intensity varies with respect to the absolute concentration and hence the ratio of fluorescence intensity of Sox2 to that of To-Pro will be the equal to the ratio of concentration of Sox2 to that of To-Pro. The normalized Sox2 concentration ratio, which is an absolute parameter, was compared across the different cell stages of early mouse embryo.

### Time laps imaging for detection of GFP signal in growing embryos in culture

Two cell-stage embryos, derived by crossing of Sox2-GFP heterozygotic females (Ellis, Fagan et al. 2004) and males, were cultured in M2 media on the Zeiss microscope (Axio Observer D1, Zeis) stage in appropriate culture conditions (CO_2_-5%, 37°C). Fluorescence images were captured in 6-hour intervals until the last blastocyst stages. GFP photo-bleaching was minimised by using a low laser power (10 μW) from 488 nm laser. The scale bar was 20 μM.

### Real time quantitative polymerase chain reaction (RT-qPCR)

All the mouse work was approved by the BRC IACUC (Biopolis). Embryos were derived by crossing of Sox2-GFP heterozygotic females (Ellis, Fagan et al. 2004) and males and collected at 3.5 dpc in M2 medium. Total RNA was extracted and purified from the whole embryos using a PicoPure RNA isolation kit (Arcturus Bioscience) and cDNA was synthesized with a high capacity cDNA archive kit (Applied Biosystems; ABI). cDNA was first pre-amplified with a pool of 48 inventoried Taqman assays (20X, Applied Biosystem) by denaturing at 95°C for 15 s, and annealing and amplification at 60°C for 4 min for 14 cycles. The pre-amplified products were diluted 5-fold and the expressions of the 48 assays were analyzed with 48/48 Dynamic Arrays on a Biomark System (Fluidigm). Ct values were calculated from the system’s software (Biomark Real-time PCR Analysis, Fluidigm). See the supplementary information for further details of methods and materials part.

## Acknowledgement

We sincerely thank, Dr Stefan Barakat and Mr Arun George Debasia for their valuable suggestions in the manuscript. We are also grateful to Dr David Rodda, Dr Kamesh Narayan and Dr Foo Yong Hwee for their technical assistance in the project. We also like to thank Ms. Elisa Hall-Ponsele for proof reading. TKM gratefully acknowledges support by a National University of Singapore graduate scholarship. Research in the IC lab was supported by the Medical Research Council of the UK. Research in the PR lab was supported by the Agency for Science Technology & Research (A*STAR). Research in the TW lab was supported by a NUS-BW (National University of Singapore / Baden-Württemberg, R-143-000-422-646) joint grant to TW and PR.

## Author Contributions

T.K.M. did experiments and wrote the manuscript, W.A. helped T.K.M in construct preparation and manuscript preparation, W.P.N helped T.K.M in FCS experiment, L.S. did immunostaining experiment, C.W. performed RT-PCR experiment, I.C. improved the manuscript by re-writing and overall work is supervised by T.W. and P.R.

## Additional Information

Please see the Supplementary Section for further information.

## Figure Legends

**Supplementary Figure S1:**
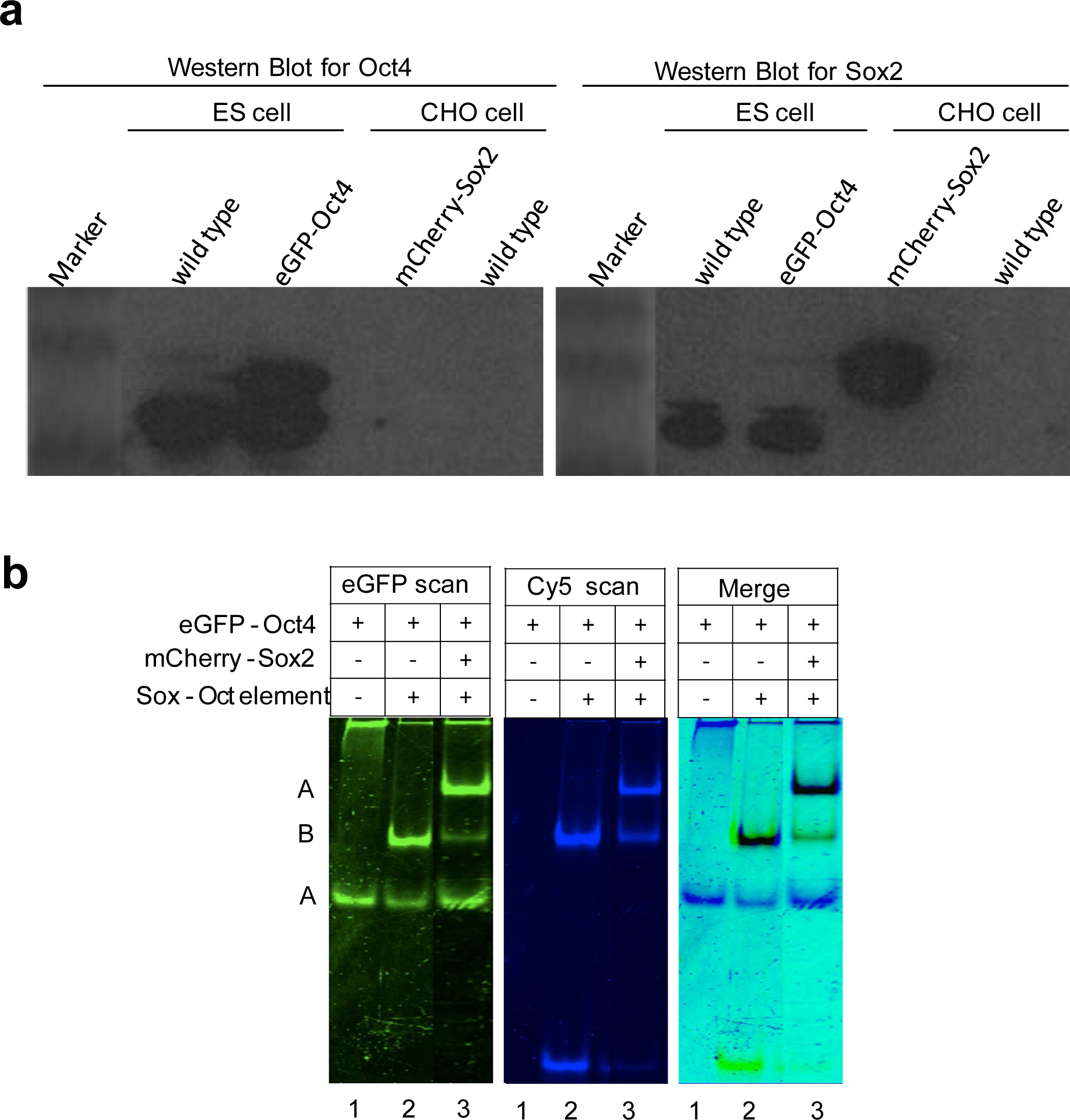
CHO cell line is ideal for fusion protein generation. (a) Western blot results for Oct4 (left) and Sox2 (right) in ES or CHO nuclear cell lysates.(b) Complex formation of GFP-Oct4 with the *Nanog* Sox/Oct motif in the presence or absence of mCherry-Sox2 by FP-EMSA.

**Supplementary Figure S2:**
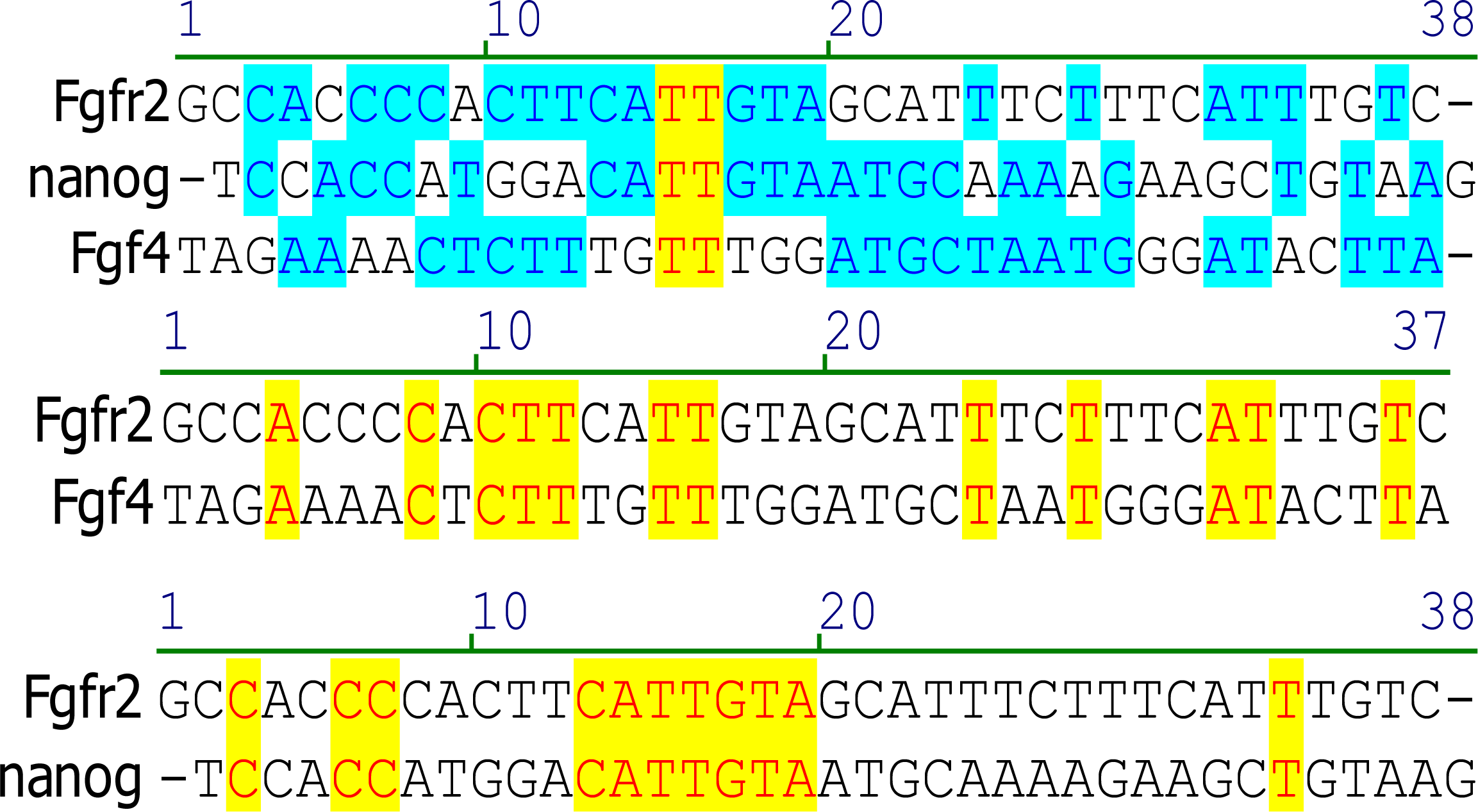
Comparison of natural DNA sequences among *Fgfr2, Fgf4*, and *Nanog* motifs. 37 bps long DNA elements containing either of *Fgfr2, Fgf4*, or *Nanog* motif were compared for all three oligos (top), *Fgfr2* and *Fgf4* (middle) and *Fgfr2* and *Nanog* (bottom). Yellow colour represents base pair similarity among variants and blue colour represents base pair similarity for less than 3 variants.

**Supplementary Figure S3:**
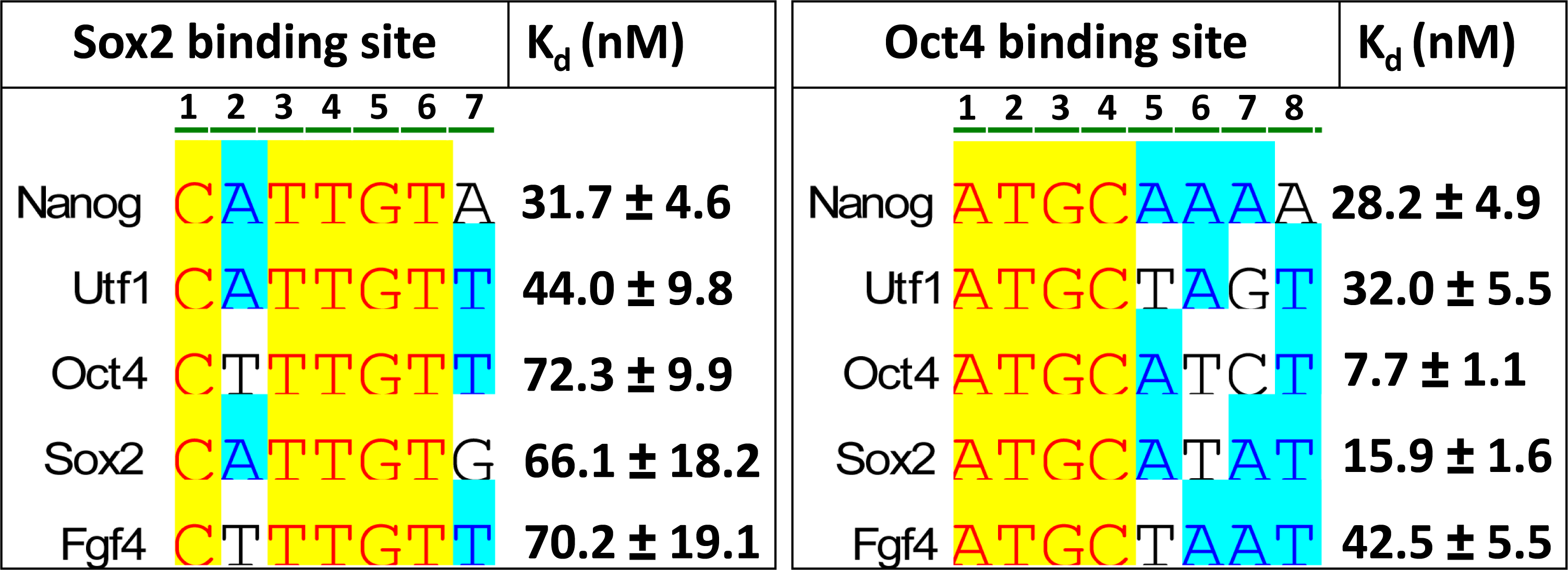
aK_d_ of Sox/Oct motifs measured for either Oct4 or Sox2. The indicated Sox/Oct motifs were compared to show the Sox2 binding site sequence (left) and the Oct4 binding site sequence (right) as well as aK_d_ values measured for either Sox2 or Oct4 alone by FCS.

**Supplementary Figure S4:**
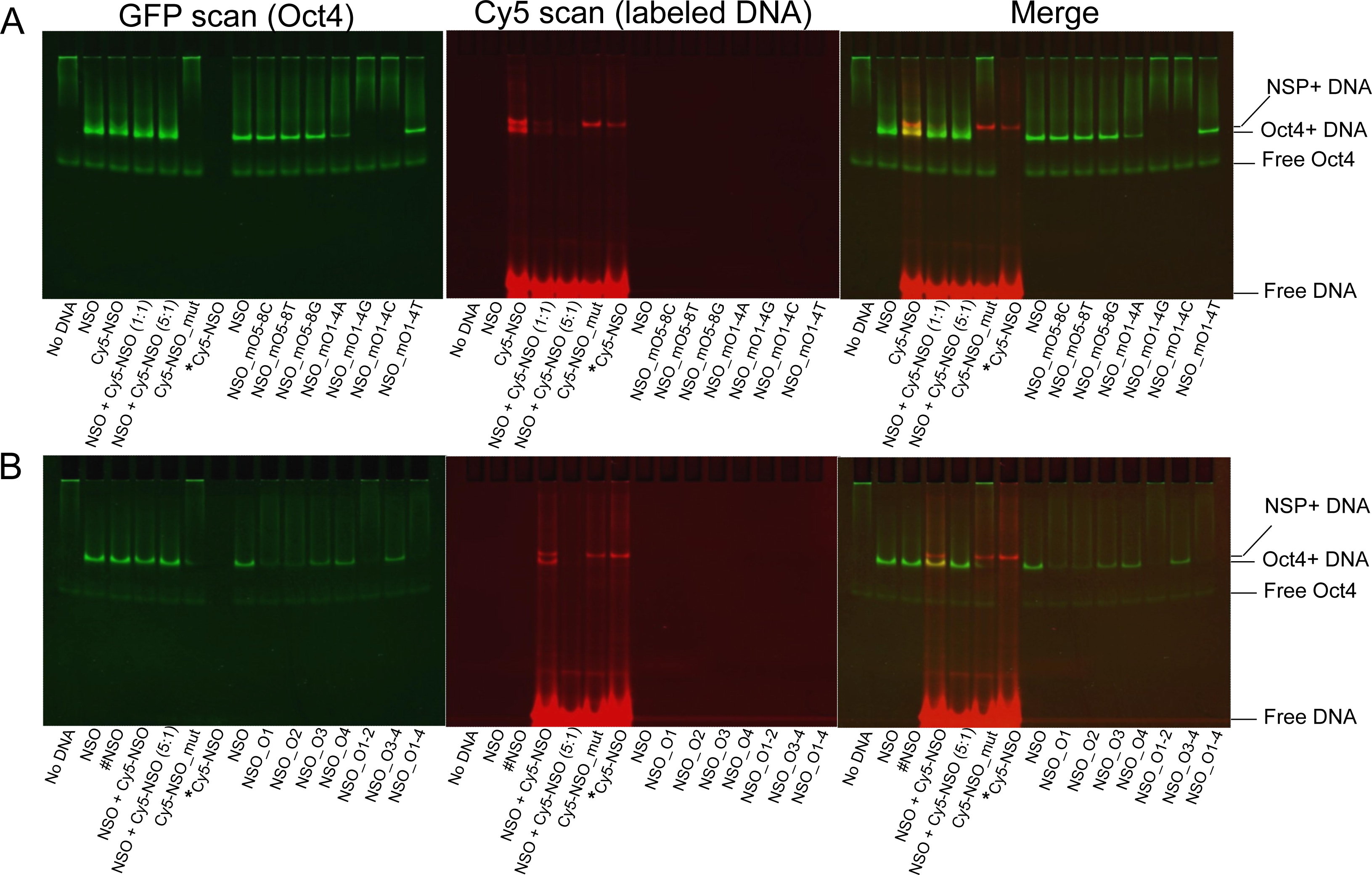
Importance of sequence specificity in Oct4-DNA binding on Nanog Sox/Oct motif. (a) The binding affinity of Oct4 to the *Nanog* Sox/Oct motif or to mutations within the octamer (ATGCAAAA) sequence is compared. Controls are shown in lanes 1 to 8 and mutation experiments in lanes 9 to 15. GFP-Oct4 was added in all lanes except lane 7 (untransfected CHO nuclear lysate). (b) The binding affinities of Oct4 to the *Nanog* Sox/Oct *cis* motif or to different mutations within the octamer (ATGC) sequence were compared. Controls were shown in lanes 1 to 8 and mutation experiments in lanes 9 to 15. GFP-Oct4 was added in all lanes except lane 7 (untransfected CHO nuclear lysate). All oligonucleotide sequences are in the supplementary information.

